# Microcapsuled entomopathogenic fungus against fire ants

**DOI:** 10.1101/505370

**Authors:** Hua-Long Qiu, Eduardo G P Fox, Dan-Yang Zhao, Jin-Zhu Xu, Chang-Sheng Qin

## Abstract

A new microencapsulation method of *Metarhizium anisopliae* based on gelatin (GE) and gum arabic (GA) is presented. Conditions to produce spheres 35-50 *μ*m relied on 1% wall material, GE:GA ratio 1:1–1:2, core:wall ratio 1:1, under pH 4.0, 35-50°C and 500-700 r/min agitation. Microcapsulation provided protection against UV and longer shelf life as compared to unencapsulated conidia. The obtained preparation proved active against the red imported fire ant *Solenopsis invicta*.

## 1. Introduction

Widespread crop use of toxic chemical insecticides has been a top concern among developing countries over the past decades (Ecobichon, 2001). Pesticides will imbalance and contaminate ecosystems, while spawning insecticidal-resistant populations (Whalon et al., 2008). The recent accelerated economic growth of People’s Republic of China had a heavy toll on the environment (WWF, 2016), and now nationals are yearning for better environmental conditions and improved safety regulation on internal goods (Song and Woo, 2008). As a result, there are solid governmental efforts to limit and/or reduce the use of chemical pesticides, such as encouraging sustainable technologies (Wang, 2017).

One established sustainable method to control insect pests is through the use of entomopathogenic fungi (Jaber and Ownley, 2017; Skinner et al., 2014). Most commonly employed fungi are *Metarhizium spp*. and *Beauveria bassiana* (Shah and Pell, 2003). Such fungi invade the integument of insect body via spores that grow hyphae into the body cavity, absorbing fluids and destroying internal organs and tissues (Roy et al., 2006). The species *M. anisopliae* can infect a wide range of insect species that is mainly employed in controlling soil pests (Krueger and Roberts, 1997; Schwarz, 1995). However, conidia preparations of *M. anisopliae* are UV-sensitive and have a short shelf life (Falvo et al., 2016; Marcos et al., 2009; Shah and Pell, 2003), which restricts large-scale production and distribution. New formulations of entomopathogenic fungi conidia are therefore an urgent necessity.

Microencapsulation is a protective process where core active materials are physically coated (Jin and Dan, 2011; Jyothi et al., 2010).

Microencapsulation methods have been extensively applied to improve pesticides, foods, enzymes, bait pheromones, where wall materials applied in a micrometric scale can effectively improve the stability of bio-active substances, as well as regulate their release (Eratte et al., 2014, Gu et al., 2010, Leefong and Cheesian, 2013, Opender Koul, 2011).

The microencapsulation formulation method known as ‘complex coacervation’ works via relatively physically gentle mechanisms of electrostatically-charged polyelectrolytes (de Kruif et al., 2004). It is particularly effective for making microcapsules of oily organic pesticides inside gelatin (GE) and gum arabic (GA) as wall materials (Eratte et al., 2014).

The literature on microencapsulation of *M. anisopliae* by complex coacervation is scarce. On a previous publication Liu and Liu (2008) tested several protocols of liquid phase coating to microencapsulate conidia of *M. anisopliae* MA126, using a range of coating materials. The authors obtained promising results (Liu and Liu, 2008), however some technical difficulties were observed, e.g. conidia did not disperse into the oil carrier, resulting in less protection to desiccation and ultraviolet radiation exposure. We herein present a novel preparation of microcapsules of *M. anisopliae* conidia obtained with an improved methodology, resulting in enhanced UV-resistance, storage life, and insecticidal activity tested against the red imported fire ant.

## 2. Materials and methods

### 2.1. Fungal culture

We isolated the highly virulent strain of *M. anisopliae* ‘M09’ from Australian soil samples (Qiu *et al*. 2014). Isolated conidia were cultured in 9-cm wide Petri dishes containing Potato-Dextrose Agar medium (200 g/L potato, 20 g/L dextrose and 20 g/L agar) held at 25 ± 2 °C and relative humidity of 75 ± 5 % for 8 days. The conidia were carefully brushed off and suspended in either 0.05% Tween-20 (i.e. controls) or dimethicone (for microcapsulation). Conidia concentrations were measured using a haemocytometer (Shanghai Qiu Jing biochemical reagent Instrument Co., Ltd.). Viability of obtained conidia was tested prior to experiments, and most of the conidia of *M. anisopliae* ‘M09’ on the surface of *S. invicta* could germinate and formed infection pegs to penetrate the ant cuticle (Fig. 1).

**Figure 1.**
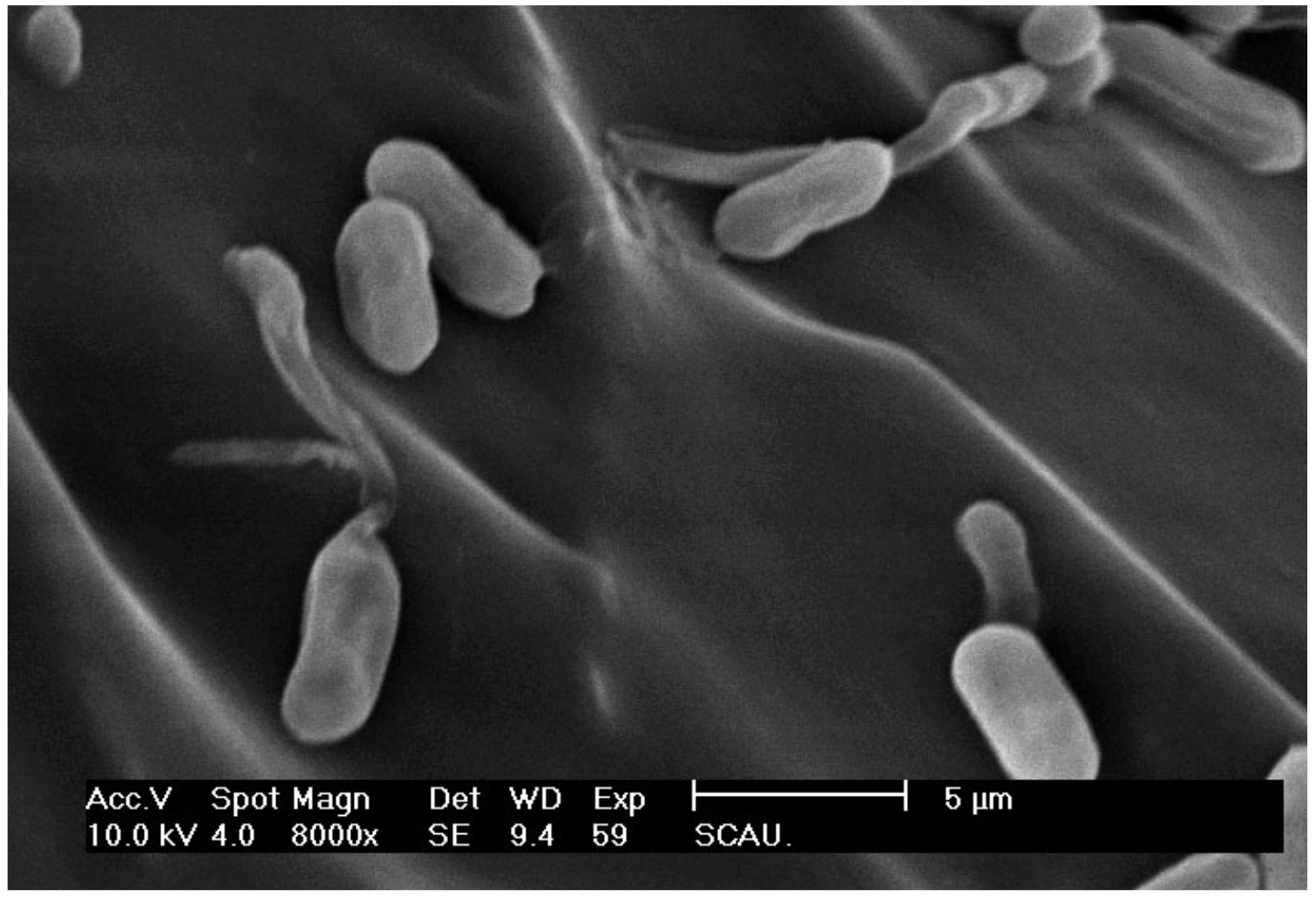
Electron micrograph of bare conidia of *Metarhizium anisopliae* M09 germinating on the thorax surface of red imported fire ants *Solenopsis invicta*, media worker, ca. 12 h following exposure.

### 2.2. General microcapsule preparation process

The employed methodology (summarized in Fig. 2) was based on the same principle of polyelectrolyte complexation as in Liu and Liu (2008), but with fundamental modifications. Background tests towards design our protocol are described in Supplementary Materials of file 1. In details, 150 mL GE (type A, with an isoelectric point between Ph 7-9) and GA solutions 2% (w/v) are prepared and mixed 1:1. The mixture is stirred for 10 min at 500 r/min until a uniform mix is obtained, and held in a 40°C water bath to prevent gel solidification. Core material (‘M09’ conidial powder) and dimethicone 1:20 (wt/wt) were then added, and gently mixed emulsified 93with a High-Shear Homogenizer at 10,000 r/min for 15 min at room temperature, towards obtaining a stable suspension of uniformly sized dimethicone micelles.

**Figure 2.**
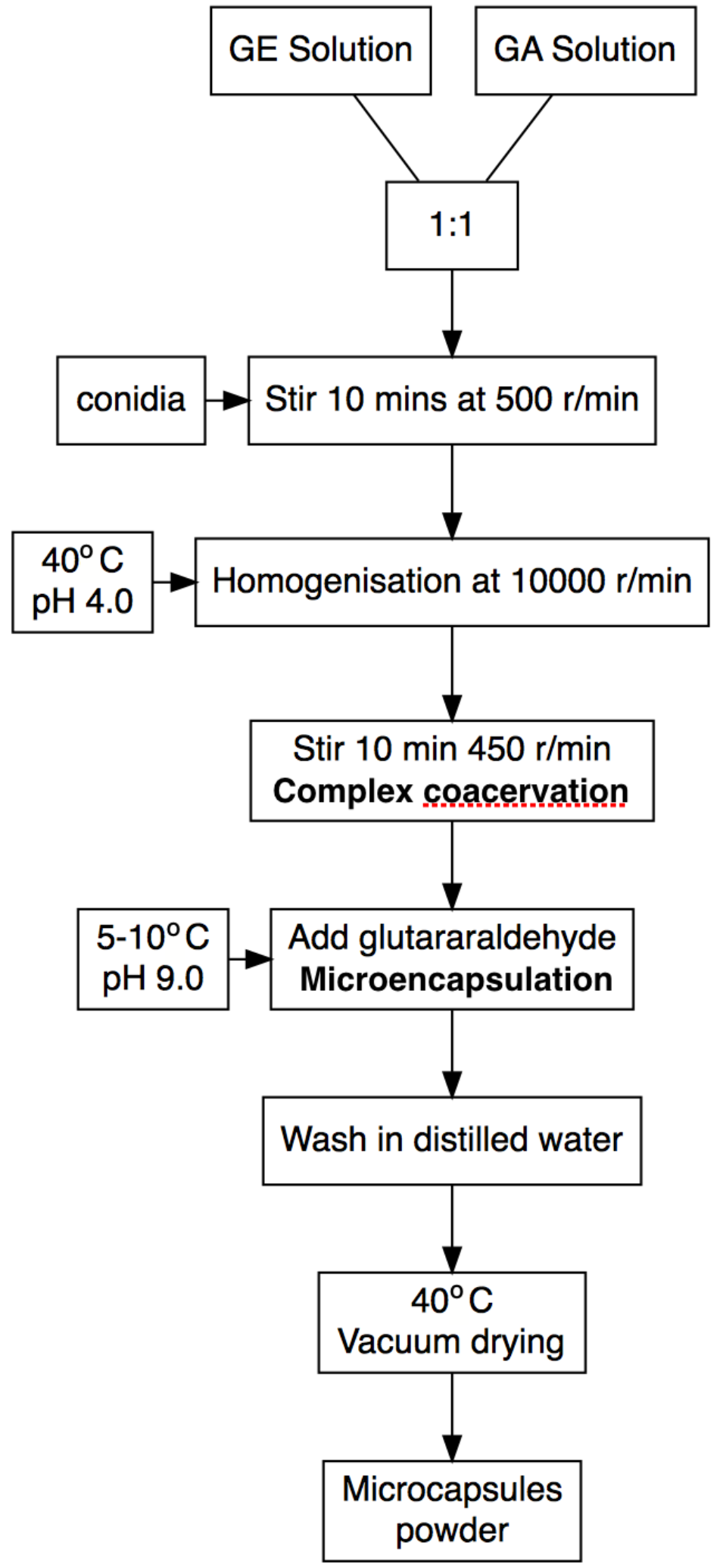
Process for preparation of Metarhizium anisopliae conidia spherical microcapsules sized 35-50 um via complex coacervation method. Preliminary tests leading to the present optimised protocol are discussed in Supplementary Materials.

This emulsion was stirred at 500 r/min in water bath temperature increased to 45 °C. As temperature stabilised the system was left stirring and pH was adjusted to 4.0 with 10% acetic, leading to complex coacervation. Coacervation was allowed to continue for 10-15 min, when temperature was lowered to 10°C. The pH was re-adjusted to 9 with 10% NaOH. Finally, wall material was fixed with 25% glutaraldehyde. Mix was centrifuged with 3000 r/min for 5 min, and supernatant was discarded. The gel microcapsules at the bottom of the centrifuge tube were washed with distilled water to remove remaining glutaraldehyde. Recovered microcapsules were held at 40°C under 0.1 MPa of vacuum in a vacuum freeze-drying machine (Shanghai Far East Pharmaceutical Machinery Co., Ltd.) until they reached a constant weight as tracked with a scales.

### 2.3. UV-susceptibility of microcapsules

Microcapsules and crude *M. anisopliae* conidia were simultaneously exposed to 15 watts UV lamps at 250 nm for 3h, and then slowly dissolved in PBS pH 7.4 for 2.5 hours, towards obtaining stable suspensions. The obtained conidial suspensions were cultured on a shaking table set to 25 ± 1°C and 120 r/min for 6 h. Germination rates of conidia were obtained to probe for UV-susceptibility of each sample. Experiment was replicated three times.

### 2.4. Shelf life of microencapsulated conidia

Microcapsules and crude conidial preparations were stored at 25 ± 1 °C and dry conditions with 30 ± 5 % RH. Samples of each preparation were taken at 0, 1, 2, and 3 months, and dissolved in PBS pH 7.4 towards obtaining conidial suspensions and cultured as above described, to obtain germination rates. Experiment was replicated three times.

### 2.5. Insecticide activity bioassay

Colonies of *S. invicta* were excavated from Guangdong Academy of Forestry, Guangzhou, China (23.200036N, 113.369204E), separated from soil as described in Banks et al. (1981), and transferred to plastic boxes (50 cm × 40 cm × 15 cm) rimmed with talcum powder. Ants were fed with *Tenebrio molitor* larvae and 25 % sucrose water every two days, and held in the dark in an incubator at 25 ± 1 °C and 80 ± 5 % humidity.

To determine toxicity microcapsules powder was mixed with sand to *S. invicta* media workers (1.15 ± 0.1 mm head width). Sand was sieved through a mesh size 18, from which 20 g of screened sand were placed in a plastic box (6 × 10 cm) with 8 ml distilled water and mixed different amounts of microcapsule powder: 0, 2, 4, 6, or 8 g. After which sixty workers of uncontrolled age were simultaneously placed in the boxes, and held in the dark incubator at 25 ± 1 °C and 85 % relative humidity. The mortality of workers was recorded once a day for seven days. Experiment was replicated three times.

### 2.6. Statistical analysis

Numeric results were analysed using RStudio v.1.1.456 using non-parametric statistics (raw data and scripts available as a Supplementary File 2). Germination rates and final mortality of ants were evaluated by Kruskal-Wallis followed by Dunn’s test for statistical significance at alpha 0.05, using R package “dunn.test”. Plots were generated with packages “reshape2” and “ggplot2”, a fluxogram was generated with “DiagrammeR”. R scripts with comments are available as a Supplementary file.

## 3. Results

### 3.1. General microcapsule preparation process

Details regarding the optimisation of the presented encapsulation protocol are discussed in Supplementary Materials of file 1. A rough estimation using the described haemocytometer was made on the number of conidia within the mixture at the beginning of the encapsulation process (ca. 3.84^10 conidia) and finally unencapsulated conidia (8.12^09 conidia), yielding an estimated % of encapsulation of 78.9% conidia.

### 3.2. UV-resistance of microcapsules

The germination rates of microencapsulated conidia were significantly higher than those of bare conidia following 3 h of UV exposure (Figure 3A; p-value: 0.0156). Germination rates were equivalent in non-exposed controls (p-value: 0.1034). Exposure to UV did not affect the germination of microencapsulated conidia (p-value: 0.3832).

**Figure 3.**
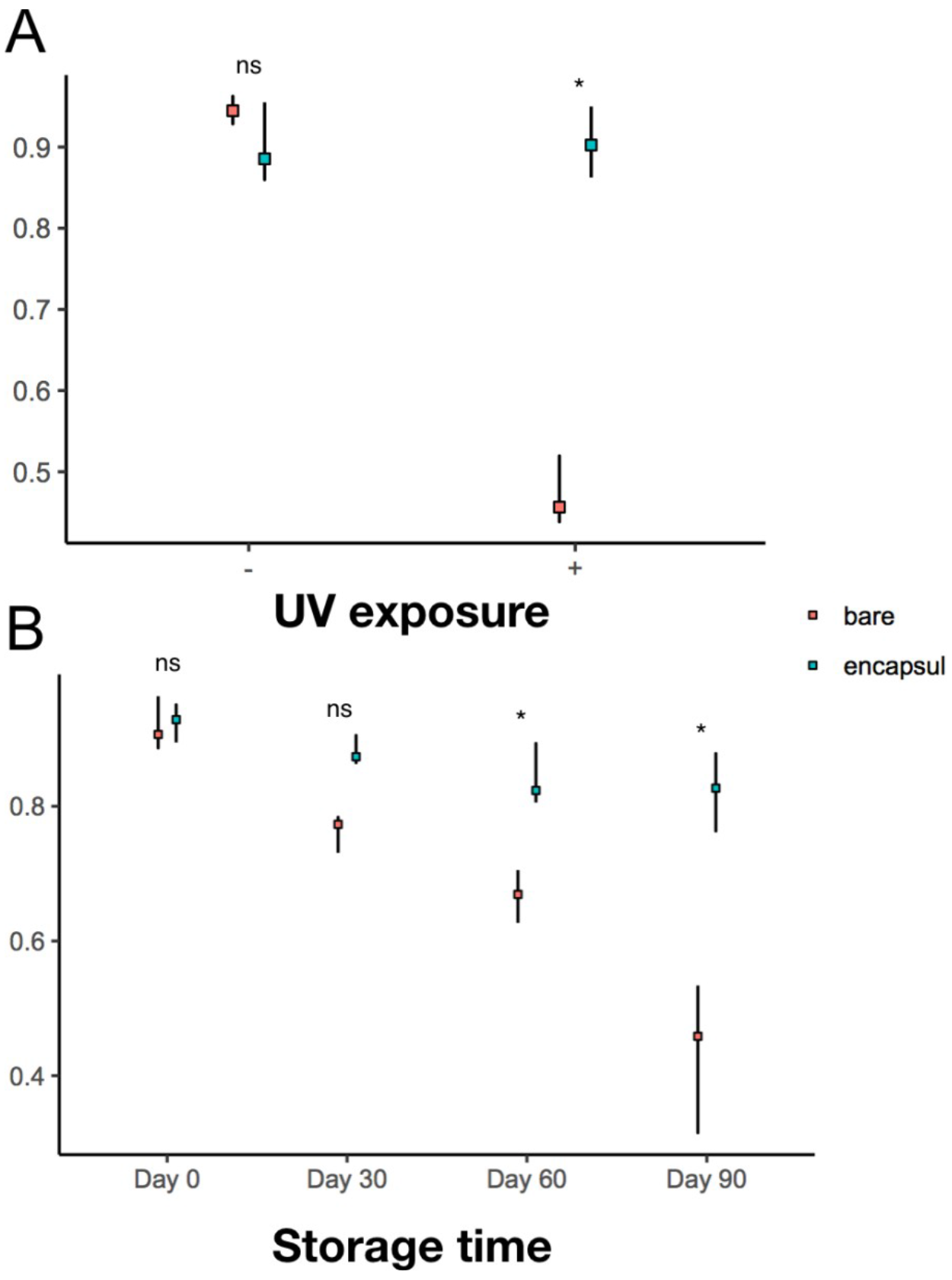
Germination rates of bare and microencapsulated conidia following (A) UV irradiation (- = no exposure; + = 2 h exposure); (B) different storage times in days 317after conidial preparation (products stored at 25oC as described in Methods). Statistics: In the plots the square □ represents the median, and vertical bars are min–max interval (n = 4). Results within each experiment were compared jointly by nonparametric Kruskal-Wallis followed by Dunn’s multiple comparison test; p-values > 0.05 were considered non-significantly (ns) different, and p-values< 0.05 were considered statistically different (*). Individual values and R plotting scripts included in Supplementary Materials.

### 3.3. Shelf life of microencapsulated conidia

Germination rates over storage months are illustrated in Figure 3B. The viability of bare conidial preparations steadily decreased over the storage period, whilst the viability of microencapsulated preparation was evidently more stable. In fact, while the mean germination rates of microencapsulated samples decreased sensibly over each month, the overall decrease was statistically negligible (refer to obtained p-values in the Supplementary File).

### 3.4. Insecticidal activity bioassay

Figure 4 illustrates the mean survival curves of *S. invicta* media workers following exposure to different amounts of microencapsulated conidial preparations. Efficiency was evaluated from final obtained rates, where the dose of 4 g was stronger than 2g (p-value: 0.0102) but not significantly different from 6 g (p-value: 0.0494) or 8 g (p-value: 0.1623).

**Figure 4.**
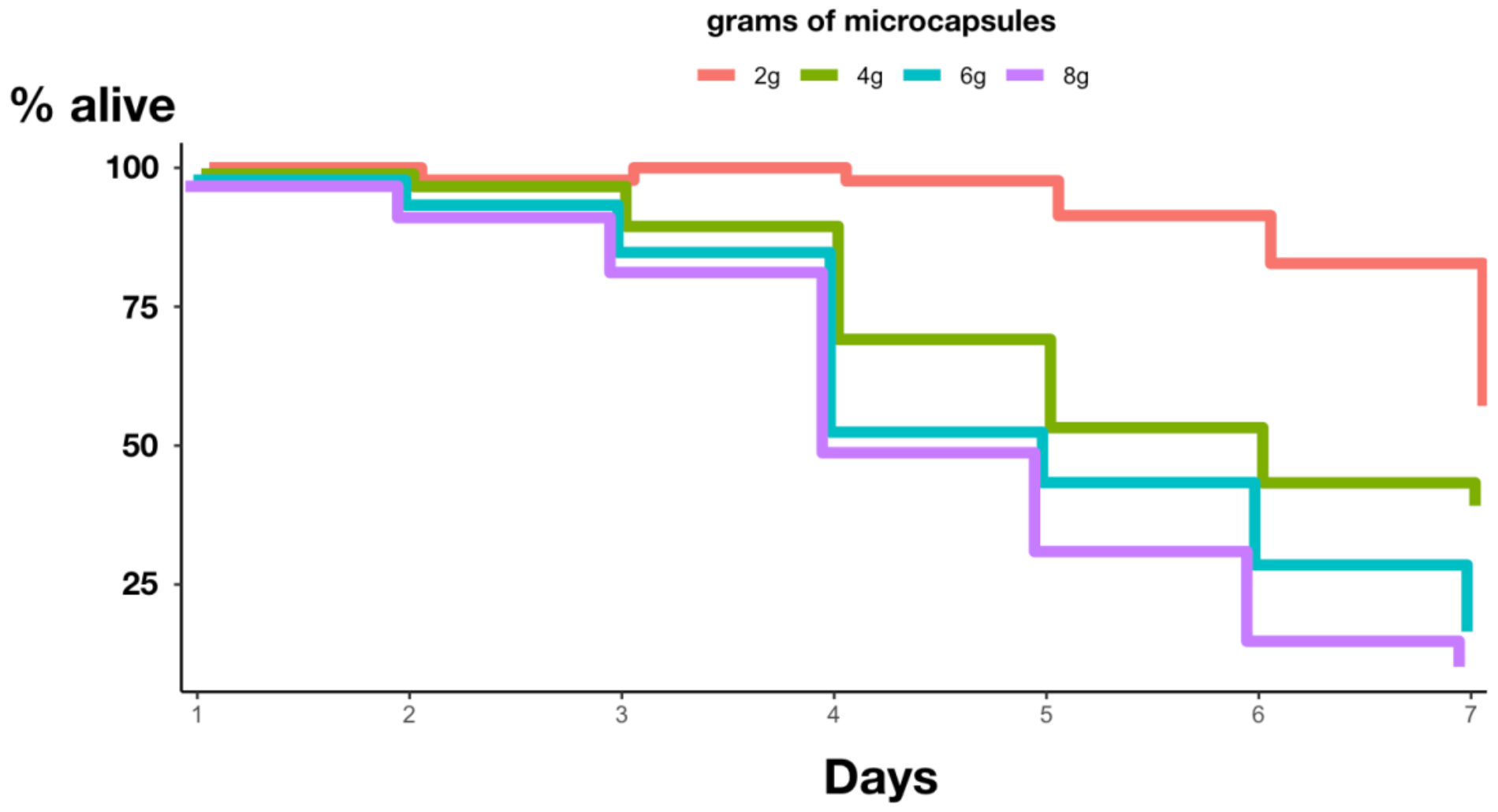
Survival curves (in days) of 30 media workers (n = 3) of the red imported fire ant *Solenopsis invicta* following exposure to different amountsof microencapsulated conidia of *Metarhizium anisopliae* prepared via the complex coacervation method depicted in Fig. 2. Individual values and R plotting scripts included in Supplementary Materials.

## 4. Discussion and conclusion

The employed M09 strain was selected for having demonstrated rapid infection against *Coptotermes formosan, S. invicta, Spodoptera litura* caterpillars in a previous investigation (Qiu et al., 2014b). The viability of the isolated sample was tested prior to present experiments (exampled in Fig. 1). Most of the conidia on the surface of *S. invicta* could germinate and formed infection pegs to penetrate the host cuticle.

Our novel method of encapsulation (Fig. 2) resulted in a conidial preparation which was more resistant than reported for preparations obtained with different protocols by Liu and Liu (2008). Compared to crude conidial preparations, the microencapsulated conidia were more stable after UV exposure and longer-lasting (Figure 3). Microencapsulated preparation also proved effectively lethal against red imported fire ants (Figure 4) at doses > 4 g per 30 worker ants.

We believe there are three possible explanations for why the process of microcapsulation was able to enhance the stress resistance of conidia. (i) Compared to other methods of microencapsulation (e.g. spray drying) complex coacervation employs gentler temperatures (Bae and Lee, 2008). Thus, likely conidia are preserved throughout the encapsulation process. (ii) The bioactive core is surrounded by a continuous layer of complexed GE:GA, isolating conidia from oxygen, UV radiation, desiccation (Mayya et al., 2003, Zhang et al., 2009). (iii) Dimethicone as a carrier works well with fungal conidia because it is atoxic and temperature-resistant (Filbry et al.,2014). Such factors taken together might be among the reasons for which the obtained microcapsules performed better than bare conidia overall. The methods described by Liu and Liu (2008) had the disadvantage that conidia did not embed into the oil emulsion, thus leaving them exposed to external stress agents.

The described microencapsulation method thus can reduce the necessary quantity of conidial required for field application from shielding conidia from environmental stress. Estimated encapsulation rates of conidia were hight (ca. 80%) but further more accurate data are needed to confirm the efficiency relative to other current methods.

## Supporting information

Supplemental Methods Details

R Script to Figures with Raw Data

