## Supplemental Methods Details for "Microcapsuled entomopathogenic fungus against fire ants"

***Optimization of complex coacervates process***

A light microscope (Olympus Deutschland GmbH, Hamburg, Germany) was used to assess the obtained microcapsules. We used uniformity in microencapulate shape and diameter as proxy to optimal conditions, where variables were: (i) GE:GA ratio, (ii) wall material concentration, (iii) core:wall ratio, (iv) pH, (v) temperature, and (vi) agitation speed. Our selected protocol (see Fig. 1 of the main manuscript) generates spherical 35-50 $\mu$m microcapsules, considered the best for microencapsulation. Herein the diameters of irregularly-shaped microcapsules was as measured from the longest width.

Parameter were tested as follows: (i) GE:GA ratios 3:1, 2:1, 1:1, or 1:2; (ii) wall material concentrations 0.5%, 1%, 2%, or 4%; (iii) core:wall ratios 1:4, 1:2, 1:1, or 2:1; (iv) pH 4.0, 4.5, 5.0, or 5.5, (v) temperatures 35°C, 40°C, 45°C, or 50°C, (vi) agitation speeds 100 r/min, 300 r/min, 500 r/min, or 700 r/min. Each factor was optimized in isolation, in sequence. Tests were replicated three times.

The influence of varied factors are illustrated in Figure S1-S6.

Concerning (i) GE:GA ratios 3:1 and 2:1 caused microcapsules to merge into large irregularly-shaped bodies (Figure S1a, b), whilst GE:GA ratios 1:1 and 1:2 led to spherically shaped microcapsules respectively 38.20 ± 319 $\mu$m and 47.6 ± 5.46 $\mu$m (Figure S1c, d).

Concerning (ii) wall material concentration 4% generates few observable microcapsules(Figure S2a) as many merged together (Figure S2b); however concentrations 1% and 0.5% led to the formation of evenly-shaped spherical microcapsules in the average diameters of respectively 36.40 ± 3.05 $\mu$m and
46.60 ± 5.03 $\mu$m (Figure S2c, d).

Regarding (iii) core-to-wall ratios,1:4 and 1:2 generated large irregularly-shaped microcapsules (Figure S3a, b) whilst evenly-shaped spherical microcapsules were obtained at ratio 1:1 (Figure S3c); no encapsulation occurs at ratio 2:1 (Figure S3d).

No microcapsules formed at (iv) pH >5 (Figure S4a), while diminutive microcapsules were produced at pH 5.0 and 4.5 (Figure S4b, c); microcapsulated within the preferable shape and sized range form at pH=4.0 (FigureS4d).

Under previous optimized conditions (v) temperatures within 35-50°C had no significant effect on microcapsules (FigureS5).

Finally (vi) stirring speeds of 100 and 300 r/min produced exceedingly large microcapsules (Figure S6a, b), while expected sizes were obtained within 500-700 r/min (Figure S6c, d).

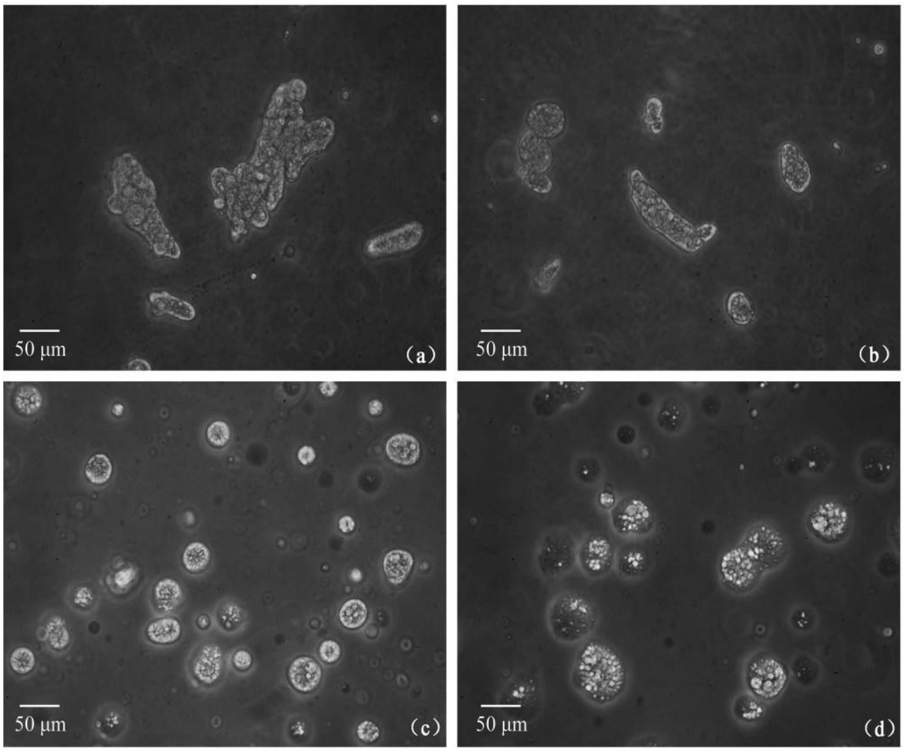


**Figure S1.** Effect of (i) GE:GA ratio (w/w) on the morphology of conidial microcapsule. (a) GE-to- GA ratio = 3:1, (b) GE-to-GA ratio = 2:1, (c) GE-to-GA ratio = 1:1, (d)GE-to-GA ratio = 1:2.

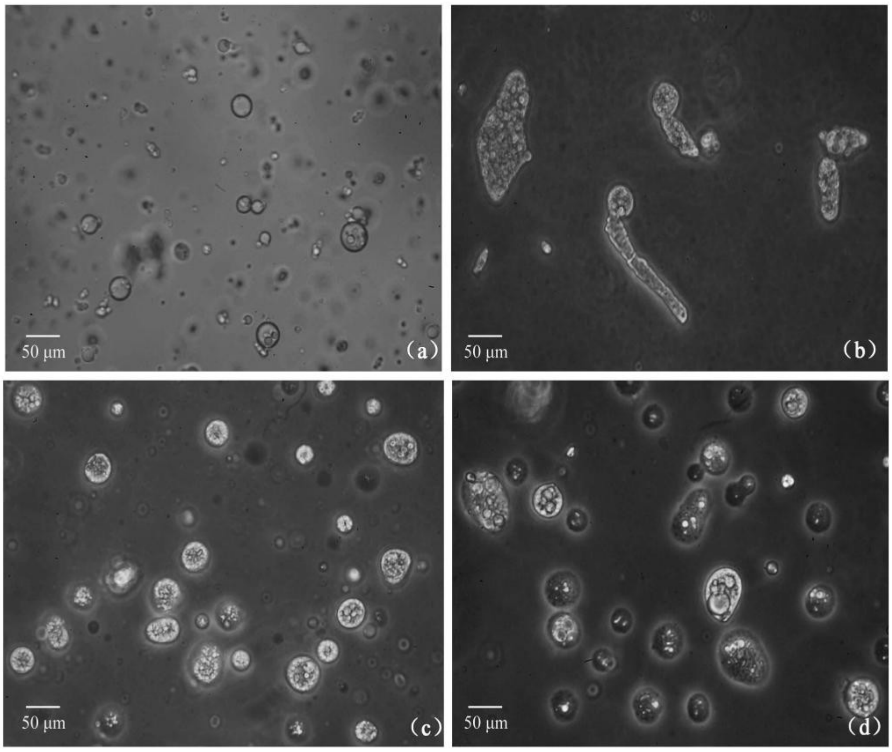


**Figure S2.** Effect of (ii) concentrations (w/v) of wall material on the morphology of conidial microcapsule. a = 4%, b = 2%, c = 1%, d = 0.5%.

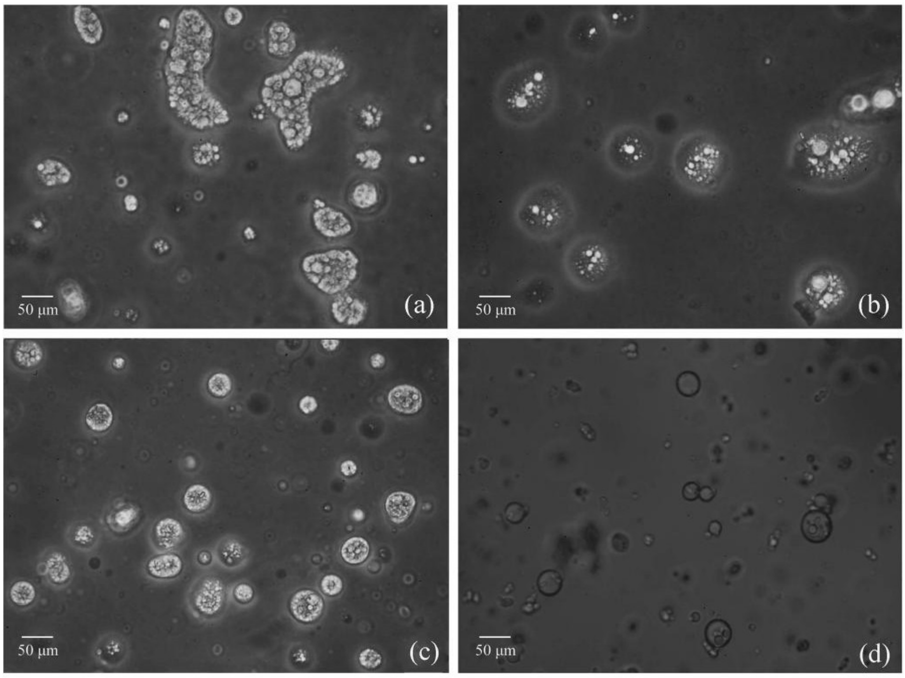


**Figure S3.** Effect of (iii) core-to-wall ratio (w/w) on the morphology of conidial microcapsule. a = 1:4, b = 1:2, c = 1:1, d = 2:1.

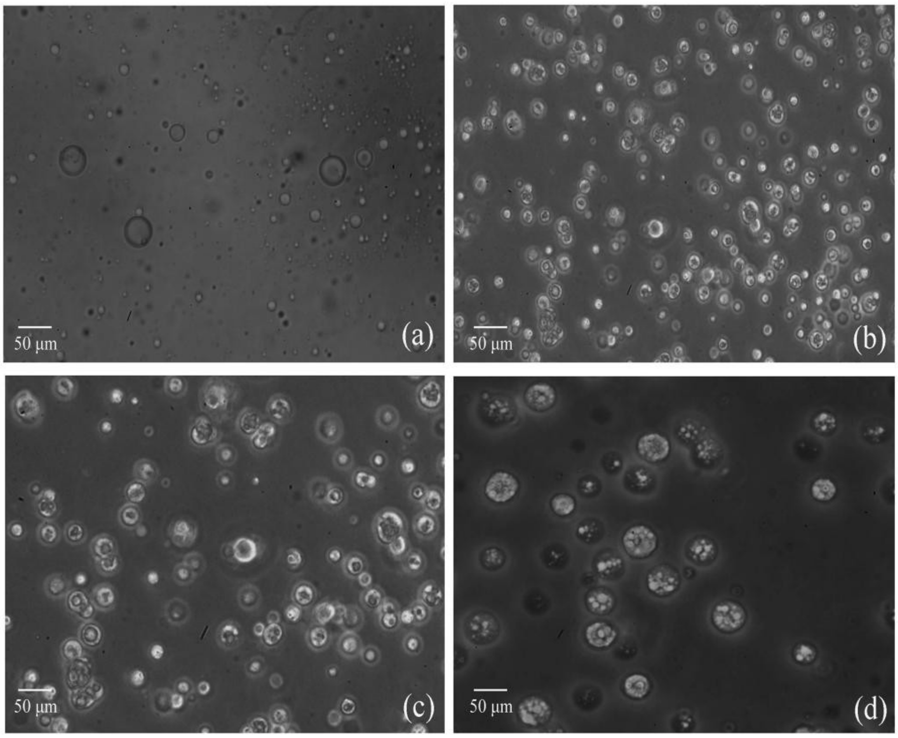


**Figure S4.** Effect of (iv) pH on the morphology of conidial microcapsule. a = 5.5, b = 5.0, c = 4.5, d = 4.0.

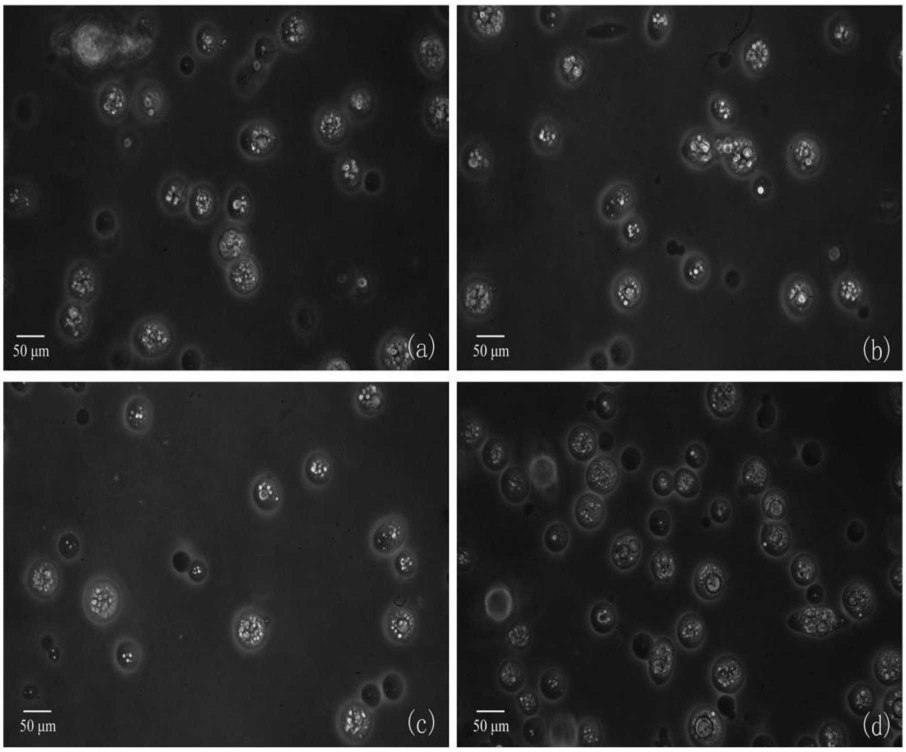


**Figure S5.** Effect of (v) temperature on the morphology of conidial microcapsule. a = 35°C, b = 40°C, c = 45°C, d = 50°C.

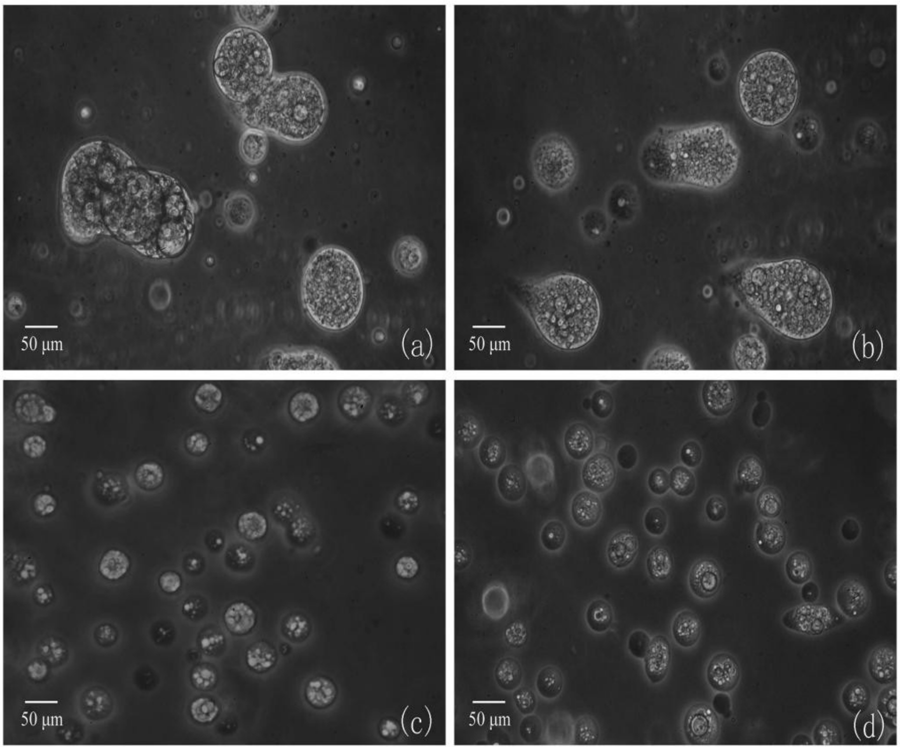


**Figure S6.** Effect of (vi) stirring speed on the morphology of conidial microcapsule. a = 100 r/min, b = 300 r/min, c = 500 r/min, d = 700 r/min.
